# Bi-phasic regulation of AIMP2 and its splice variant in PARP-1-dependent neurodegeneration

**DOI:** 10.1101/2022.04.06.487280

**Authors:** Min Hak Lee, Mi Ran Byun, Seok Won Lee, Eui Jin Lee, Young Ok Jo, Sung Hyun Kim, Wongi Seol, Kyunghwa Baek, Jin Woo Choi

## Abstract

Parthanatos is a significant molecular cause of Parkinson’s disease, in which AIMP2 aberrantly activates PARP-1 through a physical interaction. Interestingly, AIMP2 has an antagonistic splice variant, named DX2, which compromises AIMP2-induced apoptosis via p53 or inflammatory pathway. Here we suggested that DX2 binds to PARP-1 with a higher affinity than AIMP2, deactivating it and improving synaptic physiology. To deliver DX2 into deep brain areas, miR142 target sequence-embedded adeno-associated virus was designed to avoid unexpected expression in hematopoietic cells. RNAseq analysis revealed that DX2 selectively suppressed cell death-associated pathways, such as p53 and neuroinflammation. Upon a single intracranial injection, both behaviour and motility were mitigated in three animal models of Parkinsonism, induced by MPTP, rotenone, or 6-OHDA. Efficacy was observed in therapeutic model as well as preventive ones. Thus, AIMP2 and DX2 are suggested to act as an ‘ON/OFF’ switch for PARP-1. In particular, as cell survival properties of DX2 was exerted only when AIMP2 is accumulated abnormally, without its own additional tumorigenicity, DX2 could be a unique therapeutic tool for treating patients with Parkinson’s disease.

## INTRODUCTION

PARP-1 is a Janus-faced molecule participating in cell survival and death. Through polyADP-ribosylation (PARylation), PARP-1 plays a critical role in recovering damaged DNA (1). On the other hand, its overactivation causes depletion of cytosolic NAD+, forming poly-ADP-ribose (PAR), releasing mitochondrial apoptosis-inducing factor (AIF), nuclear translocation of AIF, and causing cell death (also called parthanatos) (2). Thus, while inhibition of PARP-1 can be an essential combinatorial regimen to lead synthetic lethality of DNA damaging agent such as platinum in cancer treatment, appropriate suppression of PARP-1 activation can be a therapeutic strategy to control an expansion PARP-1-induced aberrant neuronal death, such as in Parkinson’s disease and ischemia (3) (4).

Parkinson’s disease (PD) is a neurodegenerative disorder characterised by the loss of neurons in the substantia nigra pars compacta (SNpc) (5). The prevalence of PD increases with age; abnormal protein aggregates called Lewy bodies, comprising of an E3 ubiquitin ligase component called parkin, in addition to α-synuclein, are known to be formed in the brain of patients with PD. The factors that provoke PD have not been fully revealed; however, parkin mutation is known to cause autosomal recessive juvenile parkinsonism (AR-JP) (6,7). Moreover, in sporadic PD, which is a more common form of PD, parkin is commonly inactivated via S-nitrosylation, oxidative and dopaminergic stress, and phosphorylation by c-Abl (8).

Regarding the enzymatic function of parkin as a component of the SCF-like ubiquitin ligase complex, some putative substrates degraded via parkin, including aminoacyl-tRNA synthetase interacting multifunctional protein 2 (AIMP2, also known as p38 or JTV-1) (5,9), have been presented as PD therapeutic target. Since AIMP2 expression level is augmented in post-mortem brains of patients with PD, as well as in parkin knock-out mice, AIMP2 is considered to function as an important pathophysiologic factor in PD (10). AIMP2 is known to respond to genetic damage via direct interaction with p53 (11). In addition, it promotes TNF-α-dependent cell death via TRAF2, possibly acting as a multidirectional apoptotic factor (12). However, the concise mechanism by which AIMP2 affects PD has not yet been fully revealed. Recently, Lee et al. discovered that AIMP2 overexpression directly activates poly (ADP-ribose) polymerase-1 (PARP-1), and overactivation of PARP-1 causes dopaminergic neuronal cell death in AIMP2 transgenic mice (13).

AIMP2 is, however, not considered a therapeutic target for PD treatment due to its undruggability, with non-enzymatic and housekeeping scaffold properties.(14) Therefore, by suppressing the PARP-1 that is aberrantly activated by AIMP2, PARP inhibitors can be candidates for PD therapeutics. However, whether PARP inhibitors can cross the blood-brain barrier remains obscure (15). Moreover, since PARP-1 activation is important for sensing and recovery of DNA damage, long-term treatment with PARP inhibitors could induce considerable negative effects in patients with PD (16).

Although levodopa effectively alleviates PD symptoms at an early stage, it loses its effect over time, and dyskinesias or involuntary movements may remain despite levodopa treatment (17,18). To explore the possibility of a safe and specific target for treating PD, we considered AIMP2 as a target for inhibiting dopaminergic cell death. In our previous study, we had found that the splice variant of AIMP2, namely DX2, which lacks exon 2 of the full-length AIMP2, competes with AIMP2 for binding p53 (19). Contrasting features of full-length AIMP2 and DX2 were also observed in TNF-α signalling. In ovarian epithelial cells, such as SKOV3 and A2780, while AIMP2 induced apoptosis by degrading TRAF2, DX2 enhanced cell survival by activating NF-κB upon TNF-α (20).Taken together, DX2 may function as an intrinsic antagonist of full-length proapoptotic AIMP2 to balance cellular survival and death. If DX2 can block the normal function of AIMP2 in PD, dopaminergic neuronal cell death might be alleviated via DX2. This study showed that DX2 effectively inhibits the function of AIMP2 by activating PARP-1 *in vitro*.

Since CNS disease is very complex in its pathophysiology and underlying molecular mechanisms, and delivery of drugs to target cells is difficult due to the anatomical feature of CNS, more effective treatment strategies to replace existing drugs are constantly being sought (21). The present study aimed to examine the efficacy of a recombinant AAV2 vector system to effectively deliver DX2, and inhibit the function of AIMP2 in PARP-1 overactivation, both in vitro and in vivo.

## MATERIAL AND METHODS

### Cell lines and reagents

SK-SH5Y, SK-N-SH, and N2A cells were obtained from the Korean Cell Line Bank (KCLB, Seoul, KOREA). Cells were grown in RPMI-1640 supplemented with 10% foetal bovine serum (FBS) and 1% penicillin-streptomycin (HyClone, PA, USA). The transient transfection of myc-tagged or GFP-tagged AIMP2, DX2, and empty vector was achieved using lipofectamine 2000 (Invitrogen, CA, USA). PI (propidium iodide solution), MTT [3-(4,5-dimethylthiazol-2-yl)-2,5-diphenyltetra-zolium bromide], rotenone, and 6-OHDA (6-Hydroxydopamine hydrochloride) were obtained from Sigma-Aldrich (St. MO, USA). MPTP (1-methyl-4-phenyl-1,2,3,6-tetrahydropyridine) hydrochloride was purchased from Cayman Chemical Company (Ann Arbor, Michigan, USA) and dissolved in saline (0.9% NaCl, 3.75 mg/ml aliquots prepared fresh before injection). siRNA targeting DX2 (CACGUGCAGGAUUACGGGGC) was purchased from Bioneer (Seoul, Korea).

### Primary neuronal cell isolation

To prepare primary neurons, embryonic brain cortices at E18-E19 were removed and transferred to new conical tubes and then gently washed three times with PBS. The washed cortices were incubated in papain solution containing 3.5 mg/ml papain (0.5 units/g, Wako, VA, USA), 0.5 mg/ml EDTA-disodium salt (Wako, VA, USA), and deoxyribonuclease I (DNase I, 5 units/mL, Takara, Shiga, Japan) for 15 min. The digested cortices were dissociated by pipetting 12 times with a glass Pasteur pipette and filtered with a cell strainer (40-mm mesh, BD Biosciences, NJ, USA). The harvested neuronal cells were cultured in MEM containing 20% FBS and N2 supplement (Thermo Fisher Scientific, MA, USA).

### Immunoprecipitation assay

After washing the prepared cells with cold PBS, cell lysate was collected with 1% Trition x100 lysis buffer diluted in PBS after incubation at 4□ for 20 min. The 500ug of protein in supernatant was incubated with 1ug antibody for pull down for 2h and then incubated with 25ul protein A/G plus-agarose beads (sc-2003, SC; CA, USA) in 4□ for overnight. After wash the sinked bead with PBS-T (0.05% tween in PBS), 2x sample buffer was mixed with bead. For denaturation, the samples was bolied for 10min.

### western blot

The prepared protein supernatant was loaded into a 10% acrylamide gel for Western blotting electrophoresis before transferring the protein from the gel to a PVDF membrane; 5% skimmed milk solution was then used for blocking. The membrane was incubated in primary antibody diluted with Tris-Bufferd Saline (TBS) with 0.05% Tween (TBS-T) for 2 h. After three washes with TBS-T, the diluted secondary antibody was incubated for 1 h. Detection was performed after luminol (SC; CA, USA). Antibodies against c-myc (sc-40), PARP-1 (sc-8007), GFP (sc-101525) and α-tubulin (sc-5286) were purchased from santa cruz (SC, CA, USA) and tyrosine hydroxylase antibody (p40101-150) was purchased from Pel-freez (AR, USA). Poly-(ADP-ribose) antibody was purchased from cell signalling (83732S, MA, USA).

### Pulse-chase assay

HEK293 cells transfected with myc-tagged AIMP2 or DX2 were then incubated with methionine-free medium for 1 h. Then, [^35^S] methionine (50 μCi/ml) was added and incubated for 1 h. After the radioactive methionine was washed off with fresh medium, targeted proteins AIMP2 and DX2 were immunoprecipitated with the myc antibody(SC, CA US Cat. #sc-40), separated by 12% SDS-PAGE, and subjected to autoradiography using a BAS scanner (FLA-3000; Tokyo, Japan).

### MTT assay

Pre-treated or transfected SH-SY5Y cells (1×10^4) were cultured in a 96-well plate for 24h. MTT solution (10μl, 5mg/ml, Sigma) was added to each well and incubated for 3h at 37 °C. The precipitated formazan crystals were dissolved with 100μl DMSO (Duchefa) after discarding the cell culture media. Absorbance was measured at 560nm using a microplate reader (Sunrise, Tecan). The experiments were repeated three times independently.

### Primary hippocampal neuron culture and transfection

Hippocampal CA3-CA1 regions were dissected from (0–1 day-old) Sprague Dawley rats (DBL, Strain code: NTac:SD), dissociated, and plated onto poly-ornithine-coated coverslips inside a 6-mm-diameter cylinder. For live imaging of synaptic physiology, the plasmids (vGlut1-pHluorin with strep-DX2 or strep-AIMP2) were transfected 8 days after plating. Experiments were conducted 14-21 days after plating. All animal experiments were performed under the approval of the Institutional Animal Care and Use Committee of Kyung Hee University. The plasmids were incubated with 2 mM Ca^2+^, 2x HeBS (273 mM NaCl, 9.5 mM KCl, 1.4 mM Na_2_HPO_4_.7H_2_O, 15 mM D-glucose, 42 mM HEPES) and the mixture was transfected to hippocampal neurons at DIV 8 using the Ca^2+^ phosphate precipitation method.

### Optical live imaging for synaptic function

For synaptic function imaging, we have utilized pHluorin-based assay. Coverslips with neurons were mounted in a laminar-flow-perfusion system of metal chamber on the stage of a custom-built, laser-illuminated epifluorescence microscope. Live-cell Images were acquired with an Andor iXon Ultra 897 (Model #DU-897U-CS0-#BV) back-illuminated EM CCD camera with a diode-pumped OBIS 488 laser (Coherent) shuttered by TTL on/off signal from the EMCCD camera during imaging. Fluorescence excitation/emission and collection were achieved using a 40x Fluar Zeiss objective lens (1.3 NA) with 500–550 nm emission and 498 nm dichroic filters (Chroma) for pHluorin. Action potentials were caused by passing a 1 ms current pulse through platinum-iridium electrodes from an isolated current stimulator (World Precision Instruments). Neurons were perfused in saline-based Tyrode’s buffer consisting of 119 mM NaCl, 2.5 mM KCl, 2 mM CaCl_2_, 2 mM MgCl_2_, 25 mM HEPES, 30 mM glucose, 10 μM 6-cyano-7-nitroquinoxaline-2,3-dione (CNQX), and 50 μM D,L-2-amino-5-phosphonovaleric acid (AP5), adjusted to pH 7.4. All live presynaptic terminal imaging was conducted at 30 °C; all images were acquired at 2 Hz with a 50-ms exposure.

### Image analysis

Images were analysed with Image J (http://rsb.info.nih.gov/ij) using the Time Series Analyzer plugin, accessible at https://imagej.nih.gov/ij/plugins/time-series.html. Synaptic boutons of neurons were appointed as oval regions of interest (diameter, 10 pixels), and the amplitude of pHluorin-based fluorescence at synapses was counted using Origin Pro 2020. The kinetics of endocytosis was measured using a single exponential decay function.

### Rotenone-induced PD mouse model

To confirm the anti-neurodegenerative role of DX2 in vivo model, chicken-β-actin promoter induced whole body DX2 transgenic animals was provided from Dr. Sunghoon Kim (Seoul National University). the male C57Bl/6n based DX2 transgenic mice and wild type mice were used. since 8 weeks age, wild type and DX2-TG mice were treated Rotenone (Sigma-Aldrich, St. Louis, MO, USA) dissolved in 4% carboxymethylcellulose (CMC, Sigma-Aldrich) with 1.25% chloroform (Beijing Chemical Works, Beijing, China). The fresh rotenone solution was orally administrated (30 mg/kg body weight) by gavage once a day for 4 weeks. During rotenone treating, rotarod behaviour test was performed one a week and measured the latency to fall. At the last week, pole test was performed to measure the turning time above top pole and descending time from top to bottom. To confirm the dopaminergic neuron survival in substantial nigra, immunohistochemistry staining was performed with tyrosine hydroxylase antibody with cryo-sectioned brain.

### AAV production

For the production of AAV-GFP and AAV-DX2, the triple plasmid system was used. AAV-GFP and AAV-DX2 were generated by ligating oligonucleotides encoding the GFP or DX2 sequence into pSF-AAV-ITR-CMV-ITR-KanR (OXGENE, Oxford, UK) vector. AAV viral vectors were prepared and transfected with AAV2 helper (OXGENE, Oxford, UK), AAV2 Rep-Cap (OXGENE, Oxford, UK), and pSF AAV-ITR-DX2 or pSF AAV-ITR-GFP in HEK293 cells. Media and cells were then collected and AAVs were purified.

### RNA library preparation, sequencing, and data analysis

AAV-GFP or -DX2 were administrated into N2A and SK-N-SH cells. After 3 days, RNA was isolated using TRIzol Reagent (Thermo Fishern Scientific Inc., MA, USA). For control and test RNAs, the construction of the library was performed using QuantSeq 3’ mRNA-Seq Library Prep Kit (Lexogen Inc., VIE, Austria) according to the manufacturer’s orders. QuantSeq 3’ mRNA-Seq reads were aligned using Bowtie2 (22). Bowtie2 indices were either generated from genome assembly sequence or the representative transcript sequences for aligning to the genome and transcriptome. The alignment file was used for assembling transcripts, estimating their abundances and detecting differential expression of genes. Differentially expressed genes were determined based on counts from unique and multiple alignments using coverage in Bedtools (23). The RC (Read Count) data were processed based on quantile normalization method using EdgeR within R using Bioconductor (24). Gene categorition was based on searches done by DAVID (http://david.abcc.ncifcrf.gov/) and Medline databases (http://www.ncbi.nlm.nih.gov/).

### Surgical procedures

All mice experiments were performed under the Kyung Hee University Institutional Animal Care & Use Committee guidelines (KHUASP(SE)-18-101). In the 6-OHDA group, male c57bl/6n aged 8 weeks mice were injected with 4 μg/μl of 6-OHDA into the right striatum. Each mouse was ethically anesthetized using ketamine and a muscle relaxant. The anesthetized mice were then placed on the stereotaxic device with tooth and ear bars. The total volume of 6-OHDA per mouse was 3 μl. The injection was performed using a Hamilton syringe at the following coordinates: AP: +0.5 mm; ML: +1.8 mm; DV: -3.7 mm. To evaluate the efficacy of DX2 in the 6-OHDA mice model, AAV-DX2 and its control virus AAV-GFP was injected on both sides of the lesion. The injection was performed using a Hamilton syringe at the following coordinates: coordinates 1 AP: +1.2mm; ML: +1.5mm; DV: -3.5mm, coordinates 2 AP: -0.1mm; ML: +2.1mm; DV: -3.2mm. In the MPTP group, male c57bl/6n aged 8 weeks mice received 30 mg/kg of MPTP subcutaneously injected to recreate a reference model described in Petroske et al (25). All animals received a total of 10 injections consecutively for 5 days. Following the treatment, all animals were fed with sucrose solutions. To evaluate the efficacy of DX2 in the MPTP mice model, AAV-DX2 and AAV-GFP were injected into the substantia nigra. The injection was performed using a Hamilton syringe at the following coordinates: AP: -2.7 mm; ML: ±1.5 mm; DV: -4.5 mm. All stereotaxic injections were performed at a rate of 0.5 μl/min and the needle was removed out of place for another 10 min after the injection before slowly being removed.

### Drug-induced behavioral test

Every mouse in the 6-OHDA model group was behaviorally tested by using the cylinder with a total diameter of 11 cm and a height of 20 cm. All analysis was performed by a blinded experimenter to the group. Behavior tests were conducted at different points after the injection of AAV.

### Pole Test

The mouse was placed on top of a pole, and the duration of time that the mouse stayed on the pole was measured. Since mice with deficits in motor activity will fall off the pole and mice with improved motor activity will descend the pole, an increase in the time of the descent pole test indicates an improvement in motor activity.

### Apomorphine-induced rotation test

Apomorphine-induced rotation test was conducted at 2, 6, and 10 weeks post-injection of 6-OHDA. To avoid possible hypersensitization, a minimum of a 3 to 4 weeks interval was chosen. Each mouse received 0.5 mg/kg of apomorphine hydrochloride via subcutaneous injection (Sigma-Aldrich) and then was placed in the individual cylinder described above. Mice were allowed to adjust to their environment for 5-10 min before turns ipsilateral or contralateral to the lesion site. All procedures were recorded over 35 min. Results were organized by net contralateral rotations per minute.

### Rotarod test

To examine the motility of the MPTP and rotenone mouse models, a rotarod test was conducted at 3- and 15-days post-lesion. All mice were trained on the same machine with equal rotating speed to show a stable performance. Training procedures consisted of three sessions on 2 days in which each session included three individual trials, each lasting at least 200 s. All mice were trained at 5, 10, and 15 rpm (revolutions per minute). The final test was conducted at 15 rpm (three sessions, 300 s each) on the third day. To reduce fatigue, each mouse was given at least 5 min of rest between each measurement.

### Cylinder test

Forelimb movement was analysed by the cylinder test using a modified mouse version (26). Every mouse in the 6-OHDA group was tested using a cylinder (11 cm in diameter and 20 cm tall). All analyses were performed by a blinded experimenter to the group. Behavior tests were conducted at different points after AAV injection. The cylinder test was conducted at 2, 6, and 10 weeks post-injection of 6-OHDA at the onset of the dark cycle. Each mouse was placed in the plastic cylinder described above.

### Elevated body swing test

The EBS test was performed after the cylinder test in the 6-OHDA model. Each mouse was held 3 cm vertically above from the ground for 60 s. The swinging pattern of each mouse was recorded and marked whenever the mouse turned its head over 30 degrees from the axis. Each test was conducted with two examiners; one was holding the mouse while the other recorded the movement. Each test was recorded as a ratio of turns to the total number of swings.

### The SHIRPA test

For a better understanding of subtle phenotypical changes in mice, the SHIRPA (SmithKline Beecham, Harwell, Imperial college, Royal London Hospital, phenotype assessment) was performed. The SHIRPA test is a semi-quantitative analysis based on the primary method developed by Rogers *et al*. (27). Neuro-behavior assessment including tremor, grooming, spontaneous activity, locomotor activity, and clasping was measured during each set of tests. Physical phenotype assessment including palpebral closure, piloerection, body position, and tail position was measured during each set of tests. Every mouse in the MPTP model group was behaviorally tested by using the cylinder with a diameter of 11 cm and a height of 20 cm. All analyses were performed by a blinded experimenter to the group. Behavior tests were conducted at different points after AAV injection.

### Tissue preparation

At the end of the behavioral assessment, the total duration was 12 weeks in the 6-OHDA model, and 7 weeks in the MPTP model post-lesion. Mice were sacrificed after being anesthetized with isoflurane and perfused with 0.9% saline mixed with 4% paraformaldehyde (PFA, Yakuri Pure Chemicals) in phosphate buffer (PBS) at a pH of 7.4. Organs including the lungs, heart, liver, kidney, ovary, spleen, bladder, spinal cord, and brain were collected. The brains were fixed overnight in 4% PFA followed by a dehydration step with 30% sucrose in PBS. Brain samples were then frozen with optimum cutting temperature compound (O.C.T compound, Tissue-Tek) and cut on a cryotome (Leica CM1850, Leica Biosystems) into 30-μm thick coronal sections through the entire substantia nigra (SN) and the striatum. All sections were made from the posterior end of the SN to the anterior end of the striatum. The sections were stored at -20 °C in a cryoprotectant solution made with ethylene glycol, glycerol, and 0.2 M phosphate buffer.

### Immunofluorescence staining

For the staining, the floating brain tissues in cryoprotectant solution or attached cells on a cover slip were moved to slide glass. After washing with PBS three times for 5 min, blocking with 5% BSA in PBS was performed for 30 min followed by permeabilization with 0.2% triton x 100 in PBS. The primary antibody was diluted with 1% BSA in PBS, applied to the section/cells, and incubated at room temperature for 1 h. After three washes with PBS-T (0.1% tween in PBS) for 5 min, the secondary antibody diluted with 1% BSA in PBS was incubated in a dark room for 1 h. Mounting with VECTASHILD antifade mounting medium with DAPI was used for nucleus counter staining.

### Statistics analyses

The *p* values were represented as mean ± SEM and data were statistically analysed using a Student’s *t*-test or ANOVA, where appropriate. A *p* value of less than 0.05 was considered to be significantly different.

## RESULTS

### Inactivation of PARP-1 by DX2

In a previous study, it was shown that AIMP2 acts as a substrate of parkin and interacts with PARP-1; this interaction regulates neuronal cell death in PD (13). Thus, to investigate whether DX2 is a competitive inhibitor of AIMP2 and regulates neuronal cell death, we first performed a binding assay between PARP-1 and AIMP2 or DX2. We found that DX2 binds to PARP-1 more strongly than AIMP2 (Figure 1A). To study whether the expression of DX2 is important for binding of AIMP2 to PARP-1, a DX2 siRNA and vector were transfected into neuroblastoma cells, and PARP-1-AIMP2 levels were measured. The amount of binding of AIMP2 to PARP-1 decreased in DX2 overexpressing cells (Figure 1B), but in DX2 knockdown cells, the amount of binding of AIMP2 to PARP-1 is increased (Figure 1C; Figure S1A), suggesting that DX2 expression plays a critical role in the interaction between PARP-1 and AIMP2. To elucidate the mechanism by which DX2 regulates PARP-1 cleavage under conditions of oxidative stress, we firstly assessed the binding affinities of AIMP2 and DX2. When AIMP2 was expressed alone, the formation of AIMP2 homodimers was observed. However, the amount of DX2 homodimers was significantly low (Figure 1D-1E, third lane of left upper panel) compared with AIMP2-AIMP2 homodimers or AIMP2-DX2 heterodimer. Taken together, these observations suggest that DX2 prefers to bind to AIMP2 and inhibit AIMP2-induced cell death.

**Figure 1.**
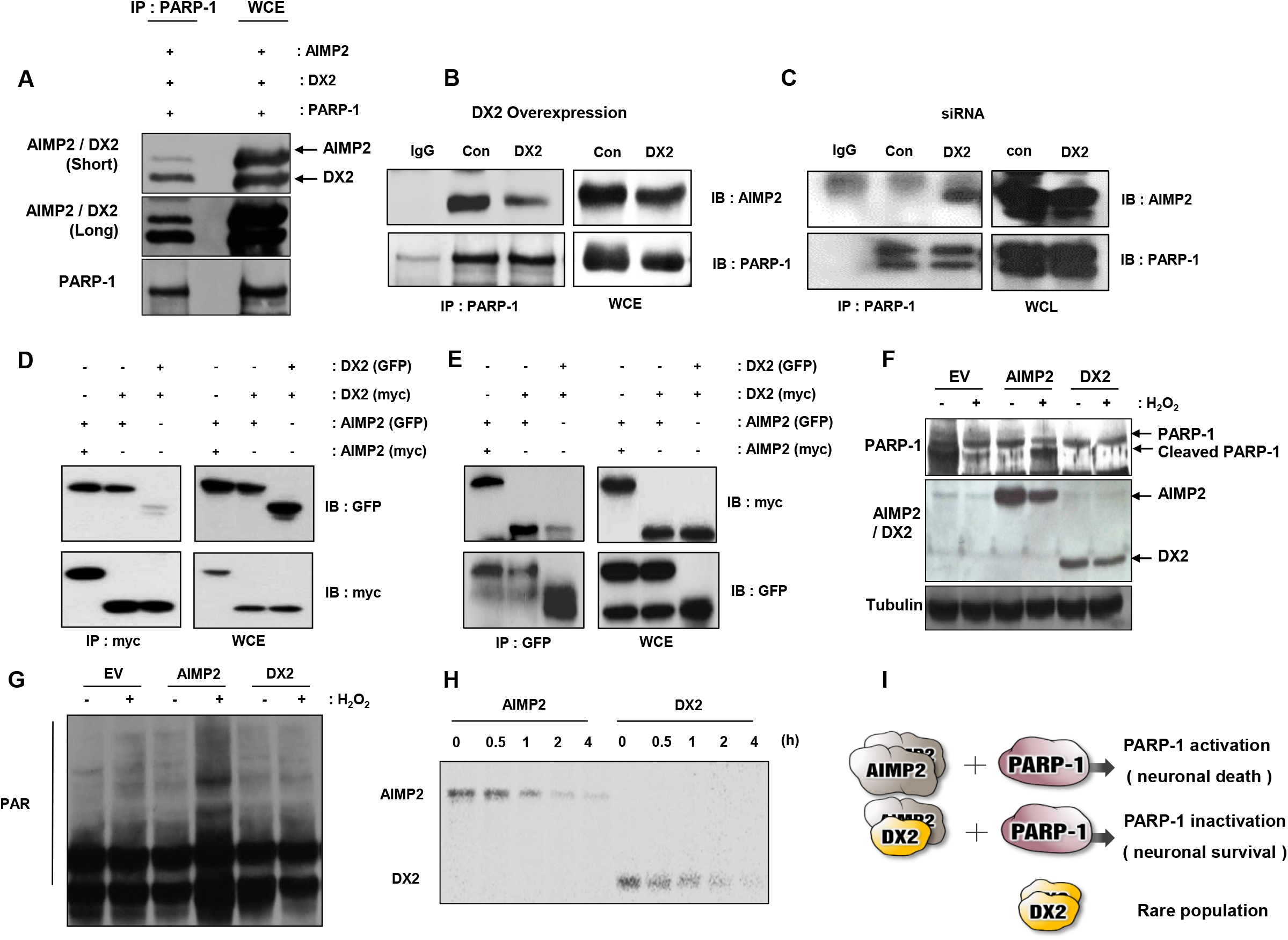
DX2 physically binds with AIMP2 to decrease PARP-1 activation. **(A)** AIMP2 and DX2 expression was induced by transfection of each plasmid in SH-SY5Y cells followed by analyses with PARP-1 pull-down assays. DX2 shows a higher affinity for PARP-1 than AIMP2. **(B)** After DX2 overexpression, binding between AIMP2 and PARP-1 was determined by Western blotting in SH-SY5Y cells. **(C)** DX2 siRNA (DX2) and control siRNA (Con) were transfected into SH-SY5Y cells and incubated for 48 h. Total cell lysates were incubated with protein agarose beads to immunoprecipitate PARP-1 bound AIMP2, which was then analysed by immunoblot analysis. **(D** and **E)** GFP-or Myc-tagged AIMP2 and/or DX2 expressing plasmid was transfected into cells and binding affinity was measured by immunoprecipitation (IP) with a myc antibody **(D)** and GFP antibody **(E). (F** and **G)** Cells were transfected with the EV (empty vector), AIMP2, and DX2; 24 h later, transfected cells were incubated with 10 μM H_2_O_2_ for 4 h. Cleaved PARP-1 levels **(F)** and PARylation **(G)** are shown. **(H)** AIMP2 and DX2 protein stability were assessed with cycloheximide-treated pulse chase assay. The samples were harvested in the time-dependent manner at 0 h to 4 h and were examined using immunoblot assays. **(H)** Schematic figure of AIMP2-induced PARP-1 activation. In the absence of DX2, AIMP2 dimers induce PARP-1 activation and neuronal death. However, in the presence of DX2, DX2 interacts with AIMP2 and inhibits PARP-1 activation.

To assess whether AIMP2 and DX2 can affect PARP-1 cleavage under oxidative stress conditions, we transfected cells with the vector expressing empty vector (EV), AIMP2, or DX2 and then treated them with hydrogen peroxide (H_2_O_2_). AIMP2-transfected cells showed significantly increased cleavage of PARP-1 when compared to the expression seen in other transfected cells under oxidative stress conditions. However, PARP-1 cleavage was not observed in DX2-transfected cells (Figure 1F). PARylation is a post-translational process, regulating biological events such as the DNA damage response and apoptosis (28,29). PARP-1 is an enzyme that recognizes damaged DNA in the nucleus, forms PAR chains, and induces degradation of damaged proteins through the PARylation. We investigated the effects of AIMP2 or DX2 on PARylation. The PARylation of AIMP2 was increased in the presence of H_2_O_2_, but the PARylation of DX2 was not altered (Figure 1G). To check the protein half-life between AIMP2 and DX2, we performed the cycloheximide treated pulse-chase assay. DX2 showed a similar stability to full length AIMP2 until 4 h from stopping of *de novo* protein synthesis (Figure 1H). Thus, these results indicate that DX2 is a critical factor for neuronal cell viability and DX2 overexpression may reduce neuronal cell death in neurodegenerative diseases. As an observation, when DX2 is insufficiently expressed, AIMP2 forms homodimers, interacts with PARP-1, activates PARP-1, and induces neuronal cell death. However, since DX2 has a significantly higher binding affinity to PARP-1 than AIMP2, the binding affinity between AIMP2 and PARP-1 was decreased in DX2 expressing cells, thereby leading to the inhibition of PARP-1 activity and the reduction in neuronal cell death. Thus, DX2 is a main regulatory protein in AIMP2-induced PARP-1 activation and cell death (Figure 1I).

### Attenuation of neurotoxin-induced neuronal apoptosis by DX2 administration

Based on these results, we suggest that DX2 is an inhibitory molecule of oxidative stress-induced PARP-1 cleavage. To check the cell viability in AIMP2 and DX2 overexpression conditions, we tested whether AIMP2 or DX2 affected cell death in the neuronal cells. An AIMP2 or DX2 expression plasmid was introduced in N2A neuroblastoma cells and MTT analysis was performed to determine the level of cell death. In normal conditions, the overexpression of AIMP2 or DX2 did not affect cell viability; however, in H_2_O_2_-treated conditions, cell death was significantly increased by the overexpression of AIMP2 and reduced by the overexpression of DX2 (Figure 2A). In the 6-OHDA treated condition, which mimics a model of PD, administration of AIMP2 induced cell death in a small but statistically significant manner compared with untreated cells. Interestingly DX2 suppressed cell death more strongly compared with H_2_O_2_-treated conditions (Figure 2B).

**Figure 2.**
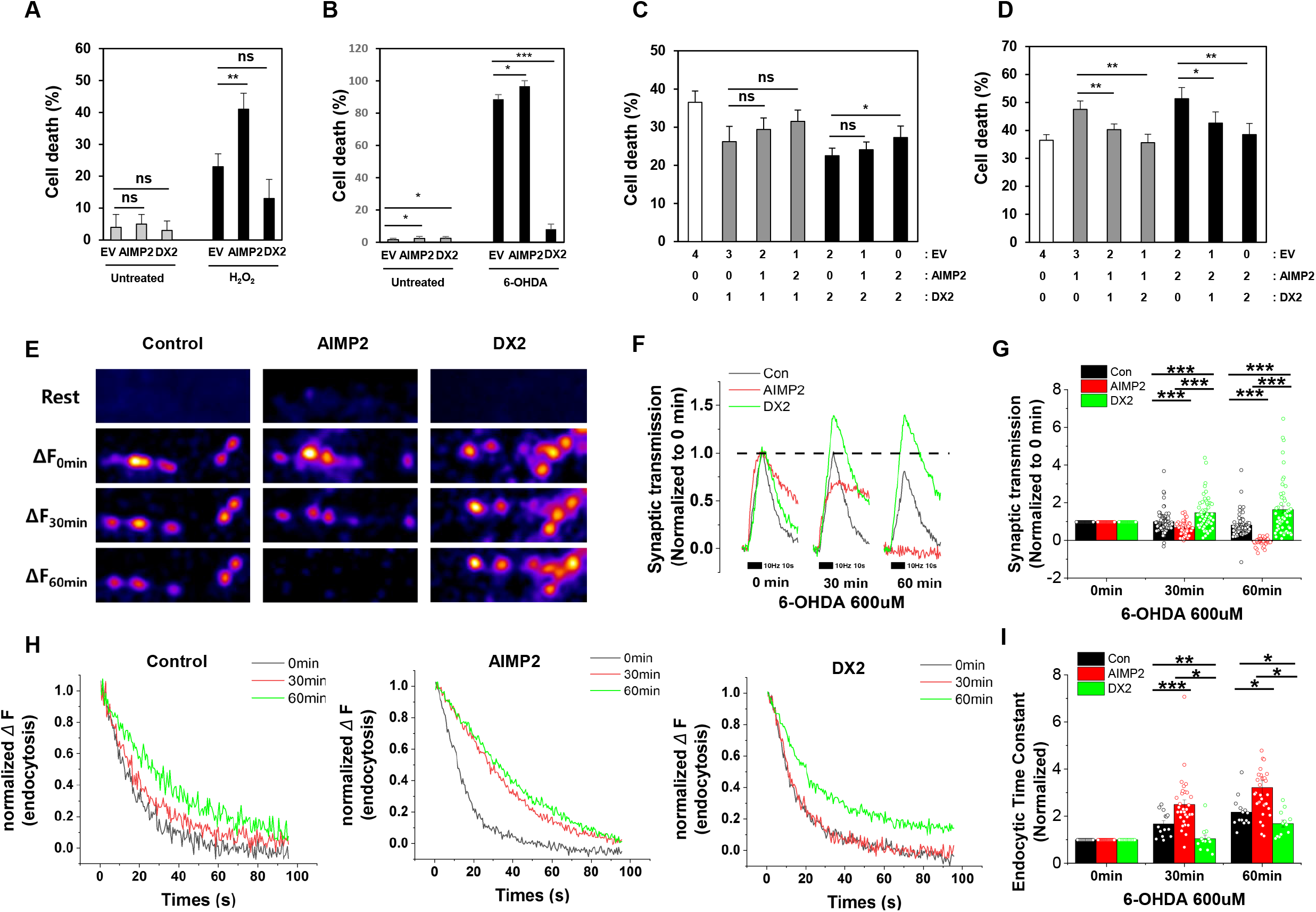
DX2 reduced neuronal apoptosis. **(A)** N2A neuroblastoma cells were transfected with empty (EV), AIMP2, and DX2 expression plasmids and incubated with or without 10μM H_2_O_2_ for 4 h. SH-SY5Y cells were co-transfected with EV, AIMP2, and DX2 expression plasmids, incubated with H_2_O_2_ for 4 h and cell viability was measured. **(B)** The cells were transfected as above and treated with 100 μM 6-OHDA for 24 h. The cell death was analysed by Propidium Iodide (PI) staining. **(C** and **D)** SH-SY5Y cells were co-transfected with EV, AIMP2, and DX2 expression plasmids, incubated with H_2_O_2_ for 4 h and cell viability was measured. The number shows the amounts of transfected DNA. **(E)** Representative images of the vGlut-pH response at synapses in Control, AIMP2-overexpressing, and DX2-overexpressing neurons. Neurons transfected with vG-pH or vG-pH with AIMP2 or vG-pH with DX2 were stimulated with 100 Action Potentials (APs) at 10 Hz in the presence of 6-OHDA (600 μM) for 0, 30, 60 minutes. **(F)** Average traces of vGlut-pH in the response of 100 APs in Control (black), AIMP2-(red), and DX2-overexpressing neurons (green) in the time course of 6-OHDA incubation as described. The traces were normalized to the peak of trace at 0 minute. **(G)** the mean values of the peak of synaptic transmission in Control (black), AIMP2-(red), and DX2-expressing neurons (green) in the time course of 6-OHDA incubation. [Control]_30min_ =0.99 ± 0.06 (n=83 boutons), [AIMP2]_30min_ =0.68 ± 0.05 (n=46 boutons), [DX2]_30min_ =1.47 ± 0.12 (n=55 boutons); [Control]_60min_ =0.81 ± 0.06 (n=83 boutons), [AIMP2]_60min_ = -0.04 ± 0.03 (n=46 boutons), [DX2]_60min_ =1.63 ± 0.19 (n=55 boutons). (***p < 0.001, **p < 0.01, *p < 0.05). Data was expressed as Means ± SEMs. **(H)** Average traces of vG-pH endocytosis after stimulation of 100 APs in Control (left), AIMP2-(middle), and DX2-overexpressing(right) neurons in the presence of 6-OHDA for 0-, 30-, 60 min. **(I)** The normalized mean values of post-stimulus endocytic time constants in Control, AIMP2-, and DX2-overexpressing neurons in the course of 6-0HDA incubation for 0-,30-, 60 min. [Con]τ_endo30min_ = 1.67 ± 0.14 (n=14 boutons), [AIMP2]τ_endo30min_ = 2.50 ± 0.20 (n=32 boutons), [DX2]τ_endo30min_ = 1.04 ± 0.15 (n=12 boutons) ; [Con]t_endo60min_ = 2.17 ± 0.17 (n=14 boutons), [AIMP2]τ_endo60min_ = 3.22 ± 0.29 (n=32 boutons), [DX2]τ_endo60min_ = 1.68 ± 0.15 (n=12 boutons). (***p < 0.001, **p < 0.01, *p < 0.05). Data was expressed as Means ± SEMs. Each endocytosis was normalized to 0 min endocytosis time constant.

Next, we studied whether DX2 or AIMP2 controls cell viability in damage-induced cell death conditions. Even when the expression of AIMP2 is increased (1 μg to 2 μg), a 1 μg transfection of DX2 suppressed AIMP2/H_2_O_2_-induced cell death to a level similar to that of 2 μg of DX2 (Figure 2C). On the other hand, AIMP2 expression level was not able to significantly affect neuronal cell death under DX2-expressing cellular conditions (1 μg to 2 μg) (Figure 2C). Therefore, DX2 expressing cells have significantly reduced cell death compared to control cells under oxidative stress conditions (Figure 2D). Taken together, the anti-apoptotic effect of DX2 seems to be much stronger than apoptosis induction by AIMP2 in the co-existant condition of the two-proteins.

Next we also reasoned whether DX2 expression has a protective effect on synaptic functionality in neurodegenerative model system. To verify this hypothesis, we utilized pHlourin-based assay which has been broadly used for monitoring synaptic functionality such as synaptic transmission and synaptic vesicle endocytosis (30,31). Primary hippocampal neurons were transfected with vGlut-pHluorin with AIMP2 or DX2, and then monitored activity-driven synaptic transmission in the presence of 6-OHDA which is a chemical inducer for PD. The treatment of 6-OHDA significantly suppressed synaptic transmission down to 80 % at 60 min compared to control neurons, and AIMP2-overexpressing neurons showed further suppression of a synaptic transmission (Figure.2E-2G). However, DX2-overexpressing neurons did not have any defect in synaptic transmission, rather facilitated synaptic function. To check endocytic activity which reflects neuronal damage, endocytosis was indirectly measured by the change of exocytic signal with the declining curve from the peak. In synaptic vesicle retrieval, while AIMP2 overexpressing condition showed lower activity than control, DX2-overexpressing neurons appeared to have an apparent protective effect in the presence of 6-OHDA (Figure 2H-2I, e.g, Compare y axis value of red curve at 30 sec), implying that DX2 supports neuronal survival against neurotoxic environments.

### Neuroprotective effect of DX2 in rotenone-induced PD animals

To study whether DX2 expression is important in PD treatment, we generated DX2-overexpressing transgenic mice, in which DX2 is overexpressed in the whole body. The transgenic mice have the same life span as wild type mice (Figure S2). We administered rotenone to control and DX2-TG mice to observe effects on parkinsonism (Figure 3A). Eight weeks after rotenone treatment, we carried out the rotarod and pole tests and analysed mice’s behavior. We observed a reduction in motor coordination in rotenone-treated wild type mice; however, an improvement in motor activity was identified in rotenone-treated DX2-TG mice (Figure 3B). In addition, we assessed the expression of tyrosine hydroxylase (TH), which is a marker for dopaminergic neurons. There was no difference in TH expression in wild type and DX2-TG mice under normal conditions, implying normal viability of dopaminergic cells in the substantia nigra pars compacta (SNpc). Under rotenone-treated conditions, TH expression was significantly reduced in wild type mice; however, TH-positive cells in DX2-TG mice showed significant recovery when compared to cells from wild type mice (Figure 3C). We also confirmed improvements in motor symptoms in DX2 overexpressing mice after rotenone treatment. In the pole test, DX2 animals turned around fater than the wild type mice (Figure 3D) and could slow the descending time assuming a strong grip over the pole (Figure 3E).

**Figure 3.**
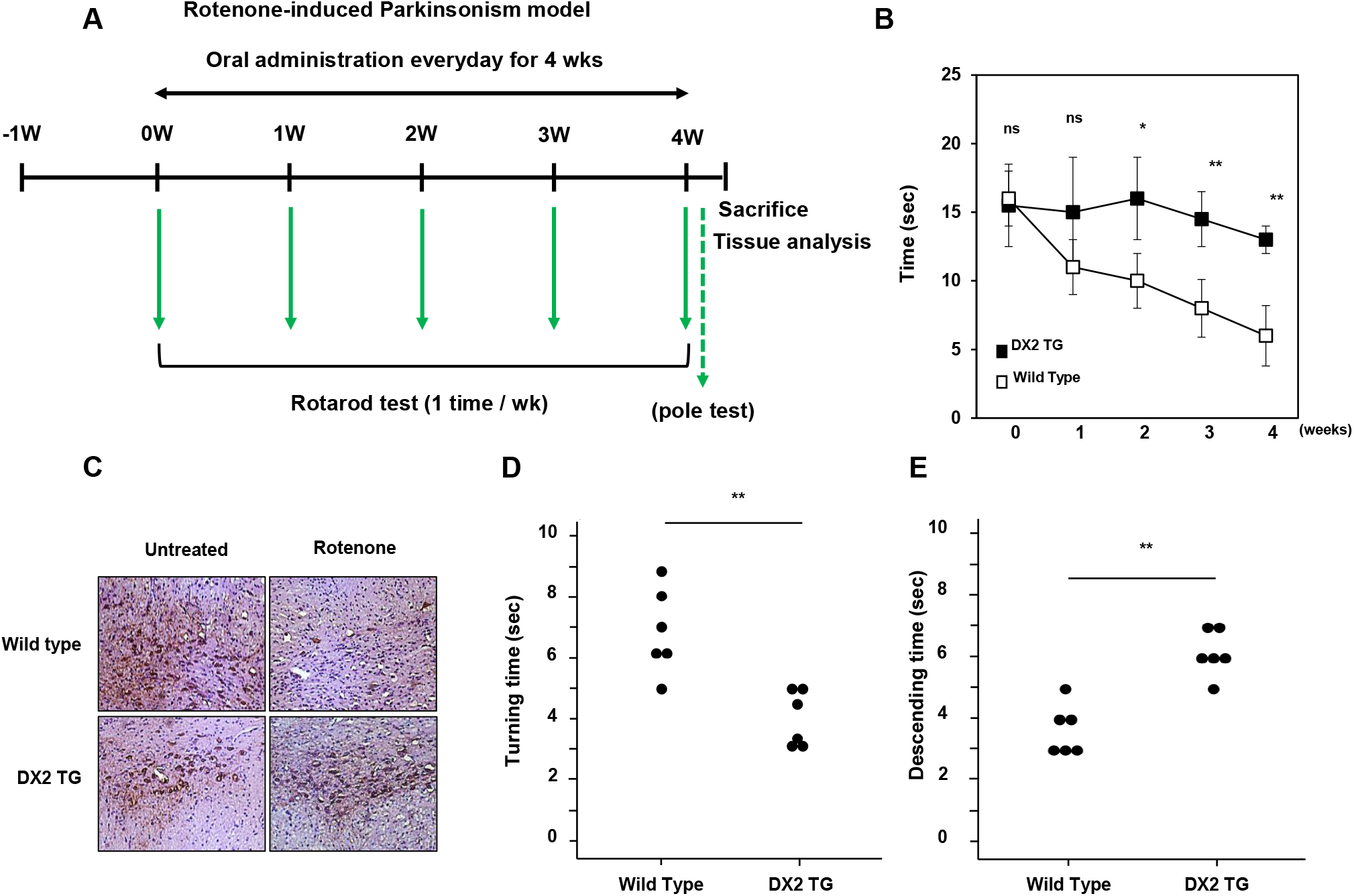
DX2 transgenic mice showed neuroprotective effect. **(A)** Schedule of Rotenone-based PD induction in DX2 transgenic mice. **(B)** Rotarod analysis. Latency to fall in rotenone-treated wild type and DX2 transgenic (TG) mice. **(C)** TH expression was analysed in the brains pf indicated mice. **(D** and **E)** The Pole test. Vertical movement and T-turn time in rotenone-treated wild type and DX2 TG mice. Animals; n=6 (in each group), *ns*; non-significant; *,*P*<0.05; **, *P*<0.01; ***, *P*<0.001.

### Design of DX2-coding self complementary AAV2

To effectively deliver DX2 to the PD patients, we introduced an adeno-associated virus serotype 2 (AAV2) delivery system. To determine which viral system to use between single-strand AAV (ssAAV) and scAAV, we found that scAAV was more effective in virus infection rates when both viruses were used to treat SH-SY5Y at different concentrations (Figure 4A and 4B). The viral vectors were injected intracranially with a minimal incision, targeting the striatum region. The expression of the transgene was tested first with the GFP signal in the control vector scAAV-GFP (Figure 4C). The expression level of AIMP2 and DX2 in the tissues was determined with Western blotting using a monoclonal antibody to detect both types. While AIMP2 seemed to be expressed broadly and stably throughout the tissues, DX2 expression showed fluctuation between tissues (Figure 4D). Immune or hematopoietic cell-rich organs such as the thymus, spleen, and colon displayed relatively high expression of DX2 compared with other tissues (Figure 4D lower bands). Interestingly, the DX2 expression level appeared to be restricted in the brain.

**Figure 4.**
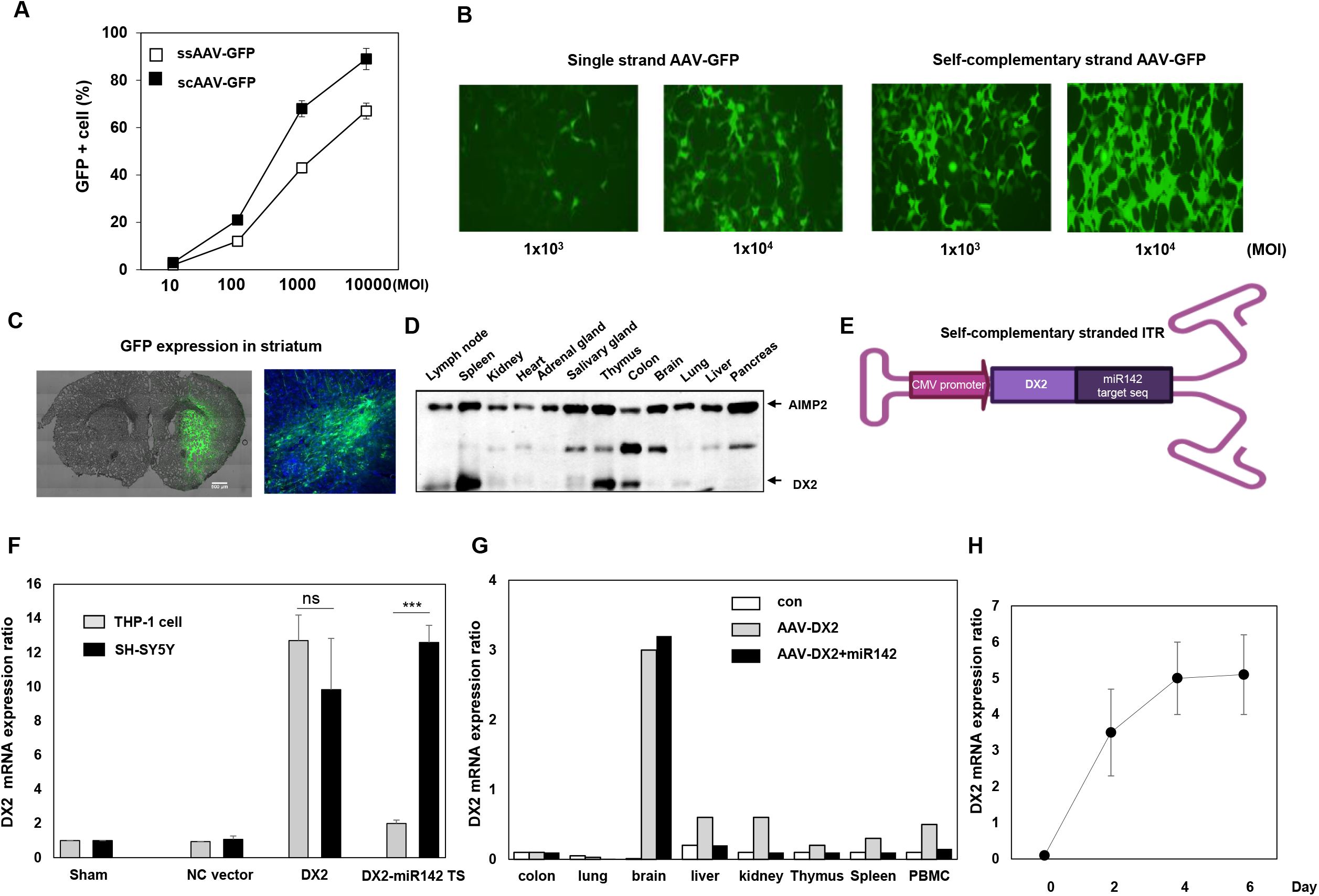
DX2 was coded in AAV2 vector. **(A)** Transduction efficacy test of AAV-GFP. The SK-SY5Y cells were infected with scAAV-GFP or ssAAV-GFP, and 48 h later, **(B)** GFP expression was observed with fluorescence microscopy. **(C)** scAAV-GFP expression in the right striatum. **(D)** Difference in AIMP2 and DX2 expression levels in mouse organs. **(E)** Scheme of scAAV-DX2 with miR142 target sequence. **(F)** Comparison of transgene expression between scAAV-DX2 and scAAV-DX2-miR142 target sequence plasmid in hematopoietic cells and neuronal cells. **(G)** DX2 expression was determined in the animal tissues at 4 days after a single intracranial injection of miR142 target sequence-inserted AAV-DX2. **(H)** DX2 expression was checked based on day after injection at the isolated substantia nigra by real-time qPCR.

Next, to confirm whether DX2 is effective as a therapeutic target for PD, we generated scAAV-DX2 (self-complementary Adeno-Associated Virus-DX2). To avoid unexpected expression in the hematopoietic cells, which showed high expression of DX2 (Figure 4D), a miRNA-142 target sequence was inserted before the 3’ ITR. (Figure 4E).

Both the AAV-DX2 viruses with or without the miR142 target sequence could infect THP-1 and SH-SY5Y cells, but only AAV-DX2 with the miR142 target sequence evaded hematopoietic cells and was only delivered in the neuroblastoma cells (Figure 4F). In AAV-DX2 without the miR142 target sequence, its mRNA level is detectable in peripheral blood monocytic cells (PBMC) and a few PBMC retaining tissues such as the spleen and kidney. Expectedly, the miR142 target sequence effectively lowered DX2 expression in untargeted tissues to the similar level with untreated animals (Figure 4G). DX2 expression showed a saturated time point at 4 days after administration with the viral vector (Figure 4H).

### Suppression of neuronal death-associated cellular signalling by AAV2-DX2 transduction

To determine whether AAV-DX2 treatment has the same effect as DX2 overexpression, we infected various cells, such as primary neurons (neuron), mouse embryonic fibroblasts (MEF), hepatocytes, MSC (Mesenchymal stem cells), and neuroblastoma cells with AAV-DX2. AAV-DX2-transduced cells (DX2) commonly showed significant decreases in cell death when compared with AAV-GFP-transduced cells (GFP) after H_2_O_2_-treatment (Figure 5A), implying that AAV-DX2 is an effective anti-apoptotic agent. Interestingly, under normal conditions, cell viability was not affected by the presence or absence of AAV-DX2. Therefore, DX2 functions only when a stressor induces apoptotic conditions (Figure 5B). We then tested the effects of AAV-DX2 in an *in vivo* mouse model. Rotenone treatment was used to induce PD and AAV-DX2 was injected into the substantia nigra. To identify possible therapeutic effects in our mouse model, we performed the vertical pole test. We observed an improvement in the behavior of AAV-DX2-transduced mice associated with the reversal of the neuronal damage caused by rotenone (Figure S5A). In addition, using the TH-stained cell technique, we confirmed that damaged dopaminergic cells in rotenone-treated mouse brains showed significant recovery in AAV-DX2-injected mice (Figure S5B). To understand DX2-related cellular pathways, RNAseq was performed. DX2 was predicted to be involved in suppression of p53-associated cell death pathway and TNF-α or interleukin-related signalling (Figure 5C-5D). The gene set was analysed based on two cell types, SK-N-SH (Figure 5E) and N2A (Figure 5F), to assess differently expressed genes (DEGs) that were categorized into signalling pathways or in ontology database. DX2 appeared to reduce the inflammatory or apoptotic signalling in both N2A (Figure 5E) and SK-N-SH cells (Figure 5F), displaying an abundancy of downregulation of related genes. The statistical significance was visualized in the distribution of a p-value graph for all of the pathways post DX2 overexpression (Figure 5G-5H)

**Figure 5.**
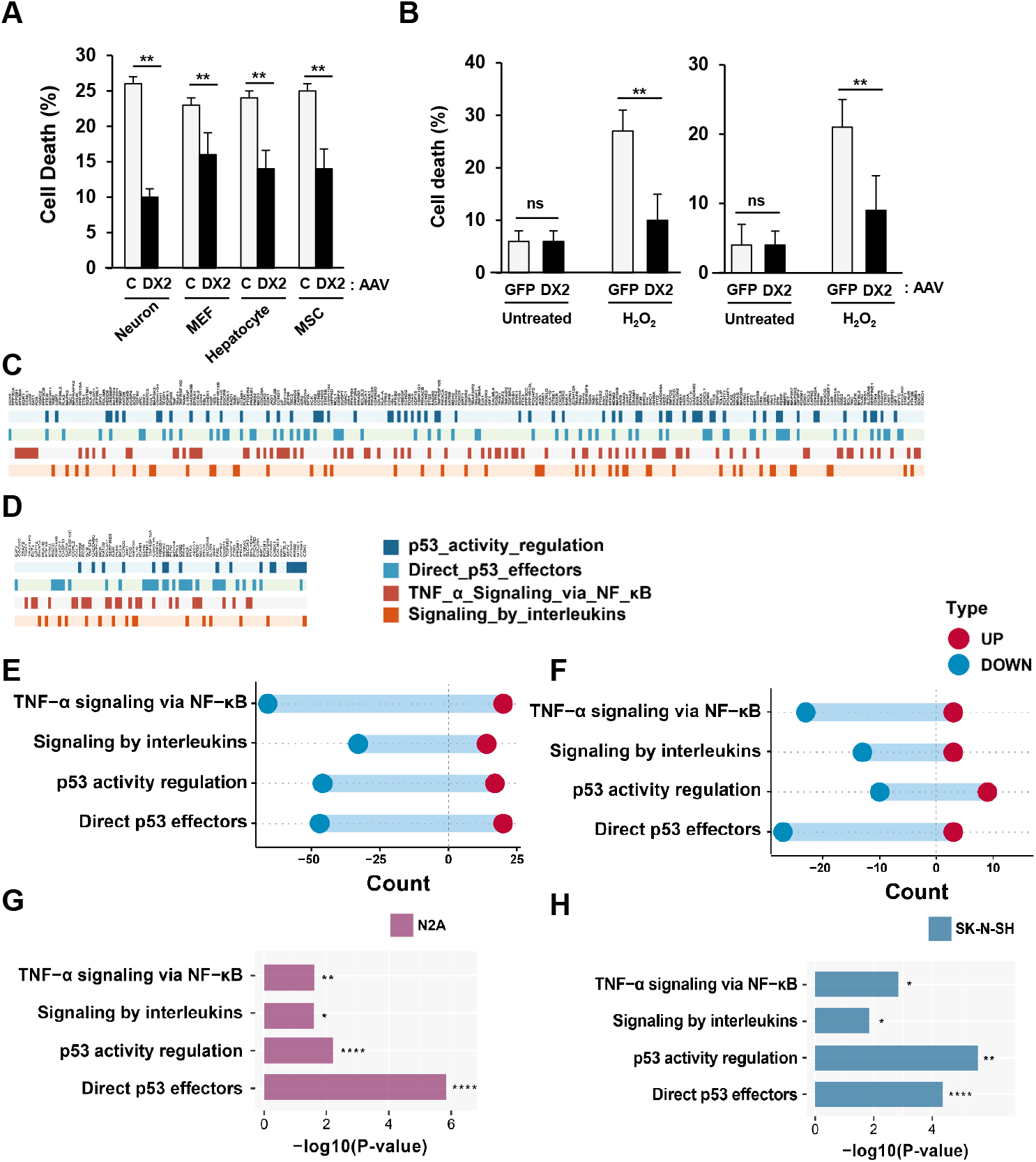
AAV-DX2 has anti-apoptotic effects and alters cellular signalling. **(A)** Cytotoxic effects in the primary neurons (Neuron), MEF (Mouse embryonic fibroblasts), hepatocytes, and MSC (Mesenchymal stem cells). After H_2_O_2_ treatment, a decreased cytotoxic effect was observed in DX2 transduced cells (DX2) when compared with control-transduced cells. **(B)** DX2 is not required for normal cell growth in SH-SY5Y (left) and primary neuronal cells (right). In oxidative stress conditions, DX2-infected cells (AAV-DX2, DX2) show decreased levels of cell death when compared to their control counterparts (AAV-GFP, GFP). **(C** and **D)** Enrichment plot of RNAseq of AAV-GFP or AAV DX2-infected neuroblastoma cells SK-N-SH and N2A. **(E** and **F)** Graph of gene counts representing cell death and inflammatory related pathways were downregulated. **(G** and **H)** p-value plot of the signalling pathway changed by DX2 overexpression. *ns*; non-significant; *,*P*<0.05; **, *P*<0.01; ****, *P*<0.0001, *t*-test.

### Neuroprotective role of DX2 in the 6-OHDA-induced PD model

To observe the effects of DX2 in the 6-OHDA-induced PD model (32), mice were first given an intracranial injection of AAV-GFP (control) or AAV-DX2 and then were injected with 12 μg of 6-OHDA into the right striatum. Two weeks later, mice behaviors were analysed twice at 4-week intervals (Figure 6A; Figure S3). As determined using the apomorphine test, motor symptoms were significantly improved in DX2-treated mice when compared to saline or GFP-treated mice (Figure 6B). In addition, to compare the neuroprotective effects between AAV-GFP and AAV2-DX2 treated mice, we performed the contralateral forepaw contact test and the elevated body swing test. In the cylinder test, both saline and GFP mice were unequally distributed to touch. However, the touch rate of the contralateral paw and ipsilateral paw was reduced in AAV-DX2 injected group (Figure 6C). In the elevated body swing (EBS) test, head swing to the right was reduced in the AAV-DX2 injected group (Figure 6D). Taken together, mice behaviors were not different in the saline or GFP-treated mice; however, recovery of 6-OHDA-induced PD symptoms was observed in the AAV-DX2-treated mice.

**Figure 6.**
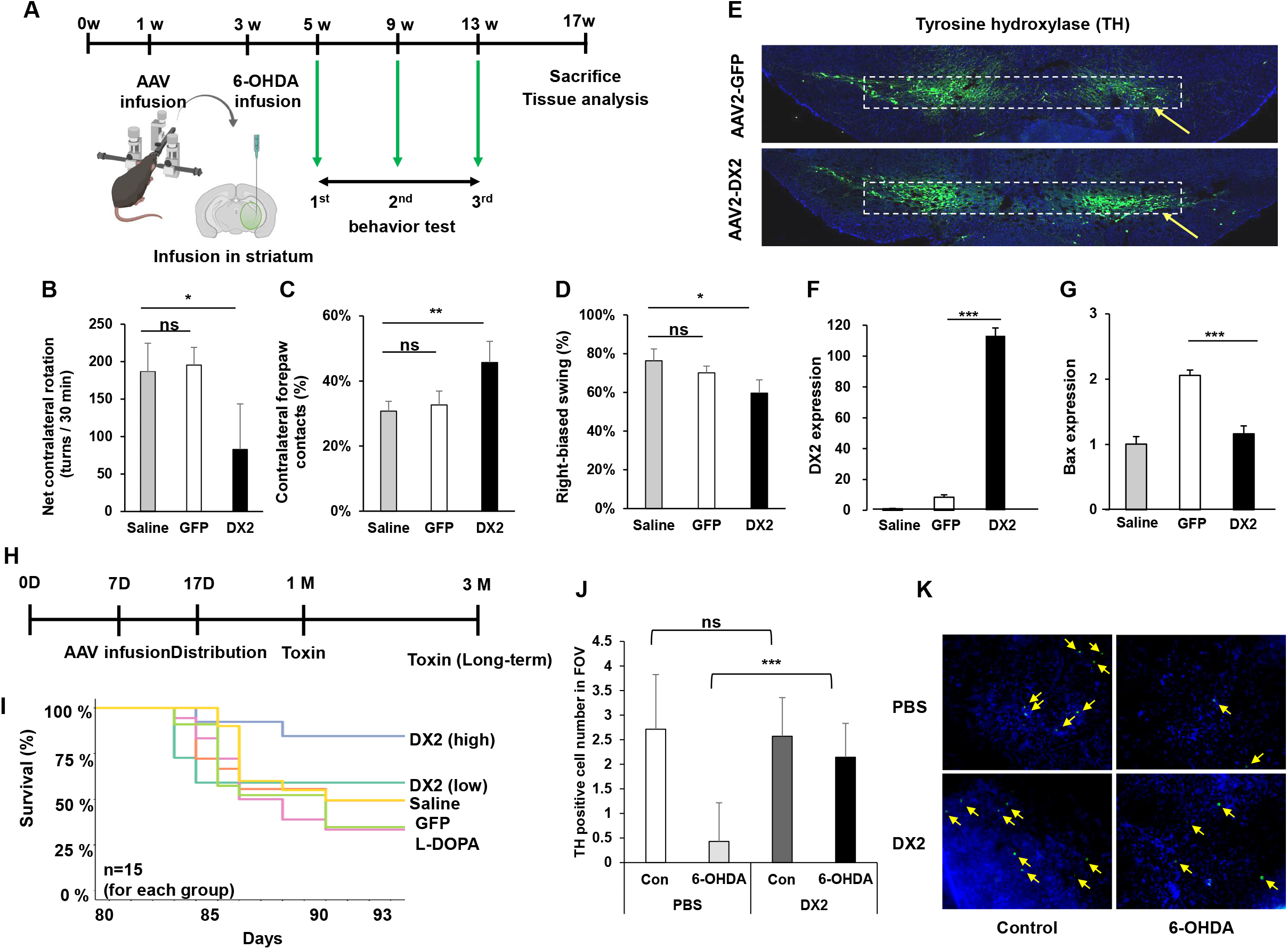
DX2 prevents behavioral deficits in the 6-OHDA-induced PD model. **(A)** Scheme of AAV-DX2 transduction in 6-OHDA-induced mouse model. **(B)** AAV-DX2-treated mouse showed lower levels of contralateral rotation compared to that of saline or vehicle (GFP), indicating that DX2 attenuated damage in dopaminergic neurons. **(C)** DX2-treated mice showed increased contralateral forepaw contacts, indicating that AAV-DX2 attenuated unilateral damage in dopaminergic neurons. **(D)** AAV-DX2 treated mouse showed less right-biased body swing. Animals; saline (saline-treated wild type mice) n=4, GFP (GFP-injected 6-OHDA-treated mice) n=5, DX2 (DX2-injected 6-OHDA-treated mice) n=11, AAV; AAV-GFP 4 × 10^9^ vg, AAV-DX2 4 × 10^9^ vg. **(E)** Immunofluorescence image of the GFP and DX2-injected mouse brain. The white square box indicates TH positive dopaminergic neuronal cells and the yellow arrows shows indicated virus injection site. **(F** and **G)** DX2 and Bax mRNA expression of naïve, 6-OHDA, and DX2-treated mice. **(H)** Scheme of the survival rate test in each mice group. **(I)** The Kaplan-Meier graph for each group. Animals; n=15, Saline indicates saline-treated wild type mice. L-DOPA, GFP, and DX2 represent L-DOPA, GFP, and DX2 injection in 6-OHDA-treated mice. AAV; AAV-GFP (GFP) 4 × 10^9^ vg, AAV-DX2 (DX2) (low) 1.6 × 10^8^ vg, AAV-DX2 (DX2) (high) 4 × 10^9^ vg. **(J)** Dopaminergic neuron counting graph. After 14 days from isolation of primary dopaminergic neurons, transduction of AAV-DX2 was performed with 1×10^4^ MOI. Before immunofluorescence, 300 μM of 6-OHDA was treated for 24 h. **(K)** Representative image of primary dopaminergic neuron staining. ns; non-significant; *,P<0.05; **, P<0.01; ***, P<0.001, t-test.

To further study the expression of TH in the AAV-transduced mice brain, we performed immunostaining. Dopaminergic neuronal cells were increased in DX2-transduced mice compare to in GFP-transduced mice (Figure 6E). Furthermore, to confirm the anti-apoptotic effect of dopaminergic neurons in DX2-injected mice, the mRNA expression level of DX2 and the apoptotic marker gene, Bax, was analysed by quantitative RT-PCR. The DX2 expression level was significantly increased in DX2-injected mice, as expected (Figure 6F), and Bax expression was reduced in these mice (Figure 6G).

The survival rate of the groups treated with saline, L-DOPA, AAV-GFP, and AAV-DX2 were analysed in 6-OHDA induced mice (Figure 6H). The survival rate was increased in DX2-treated mice (both low and high concentration) compared to L-DOPA and GFP-treated mice (Figure 6I). To confirm that AAV-DX2 prevented 6-OHDA-dependant dopaminergic neuronal cell death, transduction of AAV-DX2 was performed in primary neuron cells from wild type mice and DJ-1 null mice, a representative PD TG mouse model, before treating with 6-OHDA. The average number of dopaminergic neurons in the field of view (FOV) was decreased due to 6-OHDA, whereas the DX2 transducted group has 6-OHDA resistance in both mouse types (Figure 6K and 6L; Figure S4).

### Preventive and therapeutic effect of DX2 in the MPTP-induced PD model

In previous reports (25,33), it has been shown that MPTP produces a neurotoxin and induces severe neuronal cell death. Therefore, we also tested whether DX2 affects the MPTP-induced PD model (Figure 7A; Figure S3). MPTP induced mice to fall off the chuck at a high frequency; however, AAV-DX2 injected mice showed a reduced latency in the fall rate (Figure 7B). In the open field test, the locomotor activity of AAV-DX2 treated mice was significantly recovered compared to the naïve mice (Figure 7C). In the EBS test, the limp deficit score was increased in AAV-GFP injected mice; however, the score was significantly reduced in AAV-DX2 injected mice (Figure 7D). Taken together, significant improvements in mice behaviors were observed in AAV-DX2-injected mice when compared to AAV-GFP-injected mice (Figure 7B-7D).

**Figure 7.**
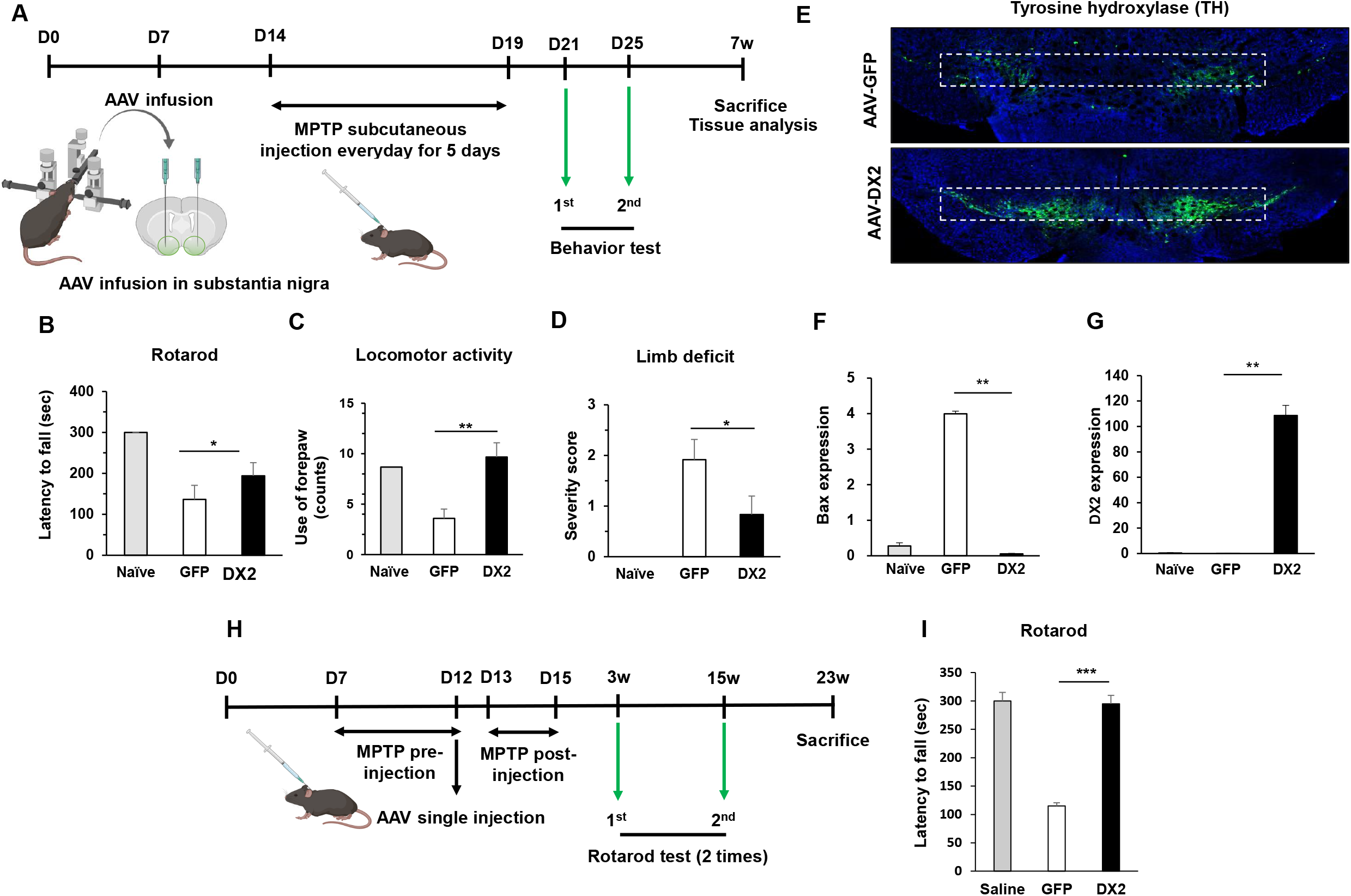
DX2 restores motor symptoms in the MPTP-induced PD model. **(A)** Scheme of AAV-DX2 transduction in MPTP-induced mouse model. AAV-DX2 was infused in substantial nigra region with stereotaxic machine and then performed subcutenous injection with MPTP for a one time per day. before measuring behavior test, training of each test was proceed. **(B)** AAV-DX2-treated mice showed slightly a longer latency to fall in the rotarod test when compared with that of vehicle (AAV-GFP, GFP) indicating that AAV-DX2 attenuated damage towards dopaminergic neurons. **(C)** DX2-treated mice showed improved locomotor activity based on the SHIRPA test. **(D)** DX2-treated mice showed a relatively lower level of limb deficit. **(E)** Immunofluorescence image of TH-positive cells in the mouse substantia nigra. **(F** and **G)** BAX **(F)** and DX2 **(G)** mRNA expression of the indicated mice brain. **(H)** Scheme of AAV-DX2 therapeutic model in MPTP-induced mouse model. before AAV injection, MPTP subcutaneous injection was proceed for a one time per day during a week. **(I)** AAV-DX2 treated mice showed recovery of movement in the rotarod test in saline-treated wild type mice, GFP-injected MPTP-treated mice, and DX2-injected MPTP-treated mice. Animals; naive n=6, GFP n=9, DX2 n=12, AAV; AAV-GFP 4 × 10^9^ vg, AAV-DX2 4 × 10^9^ vg, *ns*; non-significant; *,*P*<0.05; **, *P*<0.01; ***, *P*<0.001, *t*-test.

Similar to the immunostaining with TH of 6-OHDA treated mouse brains, TH positively stained cells were increased in DX2-injected MPTP-treated mice (Figure 7E). In addition, we observed a decrease in the expression of the apoptotic marker Bax (Figure 7F) in the DX2-transduced mice brain (Figure 7G). To investigate the therapeutic effect of AAV-DX2, MPTP was administered to mice before transduction with AAV-DX2 (Figure 7H). In the rotarod behavior test, the AAV-DX2 transduction group recovered the fall latency of the naïve group (Figure 7I). Taken together, AAV-DX2 improves behavior in PD-induced mice and has an anti-apoptotic effect on dopaminergic neurons. Furthermore, AAV-DX2 has therapeutic effect in PD.

## DISCUSSION

The most important finding of this study was that DX2, a splice variant of AIMP2 (19), can effectively turn off PARP-1 activation. The DX2 transgene encoded by rAAV2 vector system, competed with AIMP2, bound to PARP-1 more strongly than AIMP2, but did not induce its overactivation, resulting in neuronal cell survival. Over the past decade, parp-1 inhibitors have been explored in a few clinical trials to treat ischemia (34). However, it still has a technical limitation in effectively accessing the central nervous system. Since PARP-1 activation is critical for sensing and recovery of DNA damage [3], systemic or long-term treatment with PARP-1 inhibitors may cause serious side effects, like teratogenicity and anaemia (35,36).Thus, spatiotemporal or conditional inhibition would be preferred in clinical interests. DX2, a competitive antagonist of AIMP2, could be an alternative for inhibiting the PARP overactivation pathway, since AIMP2 is an upstream regulator of PARP-1 in the cell death regulation and AIMP2 accumulation results in PARP-1 overactivation and cell death, even without DNA damage (13).

Thus, using DX2 for selective inhibition of PARP-1 overactivation, only when AIMP2 accumulation is present, would be a smart strategy for treating PD effectively while avoiding unnecessary adverse effects due to the use of PARP inhibitors.

Chronic diseases, such as PD, need long-term intervention. Many allopathic medications are applicable for patients with PD, but all of them have considerable disadvantages in controlling the symptoms. Levodopa, one of the representative drugs for the treatment of PD (37,38), has many side effects, such as the end-of-dose deterioration of function or on/off oscillations, especially in patients on chronic levodopa therapy (17,18). To overcome the side effects of levodopa, many other drugs, including dopamine agonists, MAO-B inhibitors (39,40), and COMT inhibitors (41,42) are used; however, they also have other side effects. To treat patients with PD more efficiently, novel drug targets with fewer side effects than conventional drugs need to be developed; ideally, the new drug should assist the survival of dopaminergic neurons(43), besides dopamine production or secretion. In this study, DX2 was shown to improve motor activity (Figure 3, 6 and 7), rescue dopaminergic neuronal cell death (Figure 6F and 6F), and prolong life span (Figure 6J) in PD-induced mouse model, suggesting that DX2 is a critical factor for the treatment of patients with PD.

Parkin is frequently mutated or inactivated in patients with sporadic and familial PD (6,44). In previous reports, AIMP2 had been reported as a pro-apoptotic protein ubiquitinated by Parkin E3 ubiquitin ligase, and to be highly expressed in patients with PD (13). Malfunctional PD can cause AIMP2 protein to accumulate in cell, leading to aberrant cell death. A portion of the alleviated AIMP2 seems to be located in the nucleus to bind to PARP-1 (13), hence inducing parthanatos. DX2, lacking exon 2, functions as a competitive antagonist of AIMP2 (11),(12). However, since DX2 seems to act only when extra amount of AIMP2 is present, induced in a specific stress condition (Figures 1-2), logically its overexpression is non-oncogenic and it barely disturbs the tumour suppressive function of AIMP2 in normal condition (Figure 2D and 5B Figure S6) (45).

The MPTP-induced mouse model is a useful Parkinson’s disease model, since it induces acute degeneration of the nigrostriatal pathway; however, it is limited by the fact that it causes acute intoxication and does not form Lewy body (33). In the 6-OHDA-induced Parkinson’s disease model, since 6-OHDA does not cross the blood-brain barrier, it needs to be injected directly into the substantia nigra, medial forebrain bundle, or striatum to inhibit the nitrostriatal dopaminergic pathway (49). It has been reported that 6-OHDA acts specifically on monoaminergic neurons through increased ROS and quinone (50,51), and the formation of Lewy body is not induced in such mice (52). Chronic administration of rotenone causes nitrostriatal dopaminergic degeneration and Parkinson’s symptoms while increasing the α-synuclein and significantly increasing Lewy body inclusions (53). In particular, inhibition of systemic complex I by rotenone is known to induce selective effects of oxidative stress on the nitrostriatal dopamine system compared to that in the MPTP model (54). Since none of the mouse models of Parkinsonism, induced by a neurotoxin, could explain PD symptoms completely, we confirmed that the chemicals effectively induced aberrant activation of PARP-1. DX2 seemed to be an effective therapeutic tool for promoting the survival of dopaminergic neuronal cells in all the three mouse models discussed above. Further, we showed the expression level of DX2 to be important for the prevention of progressive motor deficits after neuronal damage. Collectively, we found the regulation of DX2 expression to play a critical role in improving PD symptoms, strongly suggesting that DX2 is an effective drug target for PD.

Gene therapy is a therapeutic method of directly injecting a gene to treat a genetic defect, and has been widely developed as a therapeutic option for the treatment of degenerative and intractable diseases (55). Adeno-associated viruses are currently used for gene therapy to treat PD. Clinical trials using AAV targeting PD-inducing genes, such as AAV2-hGDNF (56,57) and AAV2-hAADC (58), are currently in progress. Despite concerns about the safety of gene therapy, direct brain administration of AAV has been approved in phase I/II clinical trials of PD (59-61).

Gene therapy may be a promising strategy for accessing CNS and treating diseases, such as Parkinson’s disease, with minimal invasiveness of intracranial micro-injection. Among the various viral vectors, adeno-associated virus serotype (AAV) is accepted as a suitable gene delivery tool, owing to the following advantages: (1) long-term and stable transgene expression in non-dividing cells following the administration of a single dose and (2) relative safety of use due to the lack of pathogenicity [22]. More than 100 AAV serotypes are known to exist, of which, AAV2 is the most-selected for CNS gene therapy application due to its specific tropism, defined by CNS tissue-specific distribution and low-level systemic distribution (62). We designed a non-replicating recombinant adeno associated virus serotype 2 (rAAV2), with the CMV promoter driving the expression of transgenes DX2 and miR142-3p target sequence. scAAV system was utilised to increase the transduction efficiency. In our vector system, miR142-3p target sequence (63) was inserted to completely block the possibility of DX2 being expressed in hematopoietic cells, which form the major population of non-neuronal cells in the injected tissue area. Suppression of DX2 expression in CD45-positive hematopoietic cells was achieved by the insertion of miR142-3p target sequence, since miR142-3p is expressed only in hematopoietic cells. The miRNA 142 targeting sequence prevented DX2 expression in miR-142-3p-expressing hematopoietic cells and localised the same only to the injected tissue (Figure 4F), which is anticipated to improve safety of the therapy in clinical trials.

The current study is the first to report that overexpression of DX2 using AAV in the substantia nigra rescues motor activity and neuronal cell death in PD-induced mouse models. Considering the proven safety of AAV in clinical trials, we suggested that AAV-DX2 could be used as a therapeutic drug for patients with PD.

## DATA AVAILABILITY

All data are available in the main text or the supplementary materials.

## SUPPLEMENTARY DATA

Supplementary Data are available at NAR online.

## AUTHORS CONTRIBUTIONS

M.H.L, M.R.B, and S.W.L designed and performed the experiments. M.R.B and P.C. analysed the data. M.H.L and J.WC wrote the paper. K.B and J.W.C supervised the projects.

